# Increased bone inflammation in type 2 diabetes and obesity correlates with Wnt signaling downregulation and reduced bone strength

**DOI:** 10.1101/2024.09.26.615119

**Authors:** Giulia Leanza, Malak Faraj, Francesca Cannata, Viola Viola, Niccolò Pellegrini, Flavia Tramontana, Claudio Pedone, Gianluca Vadalà, Alessandra Piccoli, Rocky Strollo, Francesca Zalfa, Roberto Civitelli, Mauro Maccarrone, Rocco Papalia, Nicola Napoli

**Affiliations:** Department of Medicine and Surgery, Research Unit of Endocrinology and Diabetes, Università Campus Bio-Medico di Roma, Via Alvaro del Portillo 21, 00128 Roma, Italy; Operative Research Unit of Osteometabolic and Thyroid Diseases, Fondazione Policlinico Universitario Campus Bio-Medico, Via Alvaro del Portillo, 200 – 00128, Roma, Italy; Operative Research Unit of Geriatrics, Fondazione Policlinico Universitario Campus Bio-Medico, Via Alvaro del Portillo, 200 – 00128, Roma, Italy; Operative Research Unit of Orthopedic and Trauma Surgery, Fondazione Policlinico Universitario Campus Bio-Medico, Via Alvaro del Portillo, 200 – 00128, Roma, Italy; Department of Human Sciences and Promotion of the Quality of Life San Raffaele Roma Open University Via di Val Cannuta 247, 00166 Roma, Italy; Predictive Molecular Diagnostic Unit, Pathology Department, Fondazione Policlinico Universitario Campus Bio-Medico, Via Alvaro del Portillo, 200 – 00128, Roma, Italy; Microscopic and Ultrastructural Anatomy Unit, Università Campus Bio-Medico di Roma, Via Alvaro del Portillo 21, 00128 Roma, Italy; Department of Medicine, Division of Bone and Mineral Diseases. Musculoskeletal Research Center, Washington University School of Medicine, St. Louis, MO USA; Department of Biotechnological and Applied Clinical Sciences, University of L’Aquila, Via Vetoio snc, 67100 L’Aquila, Italy; European Center for Brain Research, Santa Lucia Foundation IRCCS, 00164 Roma, Italy

**Keywords:** Inflammation, Diabetes, Obesity, Bone fragility, Wnt signaling

## Abstract

Type 2 diabetes (T2D) and obesity (OB) are associated with chronic low-grade inflammation and increased fracture risk. *In vitro* studies showed that inflammation induces bone erosion and inhibits bone formation by increasing Wnt canonical pathway inhibitors. However, the impact of inflammation on Wnt pathway regulation and bone quality in T2D and OB remains unclear. To this end, we studied 63 postmenopausal women (age >65 years) undergoing hip replacement for osteoarthritis. Among these women, 19 had T2D and OB (HbA1c 6.8±0.79%; BMI 29.9±5.2 kg/m2), 17 had OB but they were normoglycemic (BMI 32.5±5.4 kg/m2), and 27 served as controls (BMI 23.1±5.5 kg/m2). Serum inflammatory cytokines by automated immunoassay (ELLA), revealed higher TNF-α (p=0.0084) and lower adiponectin (p=0.0402) in T2D, and higher IL-6 (p=0.0003) levels in OB vs controls. Gene expression analysis of trabecular bone showed increased TNF-α (p=0.0019) and SFRP5 (p=0.0084) in T2D vs controls. IL-10 was lower in both T2D (p=0.0285), and OB (p=0.0324), while adiponectin (ADIPOQ) was only lower in T2D (p=0.0041) vs controls. Interestingly, the Wnt inhibitor SOST was higher in T2D (p<0.0001) and OB (p<0.0001) vs controls. Conversely, WNT10B mRNA levels were lower in T2D (p=0.0071) and in OB (p=0.0196) vs controls, while LEF-1 were only lower in T2D (p=0.0009). WNT5A (p=0.0025) and GSK3β (p=0.0003) mRNA levels were higher only T2D vs controls. Importantly, TNF-α mRNA levels positively correlated with SOST (r=0.5121, p=0.0002), WNT5A (r=0.3227, p=0.0396) and GSK3β (r=0.3789, p=0.0146) mRNA levels, but negatively correlated with WNT10B (r=0.3844, p=0.0188) and LEF-1(r=-0.3310, p=0.0322) mRNA levels. Conversely, IL-10 was negatively correlated with SOST mRNA levels (r=0.3100, p=0.0457). ADIPOQ was negatively correlated with SOST (r=-0.3864, p=0.0105) and WNT5A (r=-0.3025, p=0.0515) mRNA levels. Moreover, SFRP5 was negatively correlated with LEF-1 mRNA levels (r=0.3991, p=0.0131). Finally, serum levels of TNF-α (r=-0.3473, p=0.0352) and IL-6 (r=-0.3777, p=0.0302) negatively correlated with Young’s Modulus, an index of bone strength. These findings suggest that increased inflammation in bone of subjects with T2D and obesity is negatively associated with Wnt pathway and bone strength, shedding light on pathophysiology of bone impairment in T2D and obesity.

## Introduction

Type 2 diabetes (T2D) and obesity (OB) are metabolic diseases characterized by systemic chronic inflammation [1]. In particular, serum levels of pro-inflammatory cytokines IL-6 and TNF-α are higher in T2D and OB [2,3]. This chronic inflammatory state contributes to tissue inflammation in insulin target tissues, and it is associated to long-term complications, including bone fragility [4]. Subjects with T2D or OB have an increased risk for fractures despite normal or higher bone mineral density (BMD), suggesting a bone quality impairment. Furthermore, subjects with elevated pro-inflammatory cytokines have been linked to an increased risk of fracture [5–7]. However, data on the direct effect of inflammation on bone fragility in subjects with T2D or OB are lacking. *In vitro* studies showed that IL-6 is mainly involved in promoting osteoclastogenesis [8]. Moreover, IL-6 contributed to bone loss *in vitro*, affecting bone marrow mesenchymal stem cells (BMCs) differentiation and cellular senescence in a model of high fat diet-induced obesity [9]. On the other hand, blocking TNF-α in bone-lining forming cells increased cell number and promoted bone formation in a model of T2D [10]. Consistently, Sun M. et al. demonstrated that subjects with T2D and fractures had higher TNF-α levels [3]. Previous data showed that TNF-α promoted bone erosion and inhibited bone formation by increasing the main inhibitors of Wnt canonical pathway [11,12]. Wnt pathway is one of the main regulators of bone metabolism. We previously showed that canonical Wnt pathway is downregulated in T2D, and it is associated with reduced bone formation markers [13] and reduced bone strength [14]. Thus, we hypothesized that increased inflammation may affect bone quality by Wnt canonical pathway in bone of postmenopausal women with T2D or obesity. Results confirmed that subjects with T2D and OB have increased bone inflammation associated with Wnt beta/catenin pathway downregulation and reduced bone strength.

## Results

### Subject characteristics

Clinical characteristics of study subjects are reported in Table 1. Age and Menopausal age were not different between groups while BMI was higher in T2D and OB compared to controls (T2D: 29.98±5.77 kg/m^2^; OB: 32.84±2.94 kg/m^2^; CTRL: 23.11±2.24 kg/m^2^, p<0.0001). As expected, fasting glucose was significantly higher in T2D compared to either OB or controls [T2D: 121.8±20.59 mg/dl; OB: 101.7±10.88 mg/dl; CTRL: 94.98±8.45 mg/dl, p<0.0001). T2D subjects had a mean HbA1c of 6.87±0.79% and a mean disease duration of 13.75±9.54 years. Diabetes medications included monotherapy with metformin (n=15) and combination therapy with metformin plus insulin and glinide (n=3). There were no differences in serum calcium, eGFR (CKD-EPI equation) and serum blood urea nitrogen, as well as in cholesterol values and triglycerides.

**Table 1.** Clinical characteristics of the study subjects Results were analyzed using unpaired Ordinary One-Way ANOVA and Tukey’s Multiple Comparisons test. Results are presented as mean ± standard deviation.

|  | **T2D (A)**<br>**n=19** | **OB (B)**<br>**n=17** | **CTRL (C)**<br>**N=27** | **P value** | **Multiple**<br>**Comparisons** |
| --- | --- | --- | --- | --- | --- |
| Age (years) | 74.90±7.22 | 71.84±9.63 | 71.92±8.36 | 0.375 | NS |
| BMI (kg/m<sup>2</sup>) | 29.98±5.77 | 32.84±2.93 | 23.11±2.24 | <0.0001 | A-B=0.010<br>A-C<0.0001<br>B-C<0.0001 |
| Menopausal age (years) | 48.65±5.18 | 49.03±4.78 | 51.13±3.47 | 0.098 | NS |
| Fasting glucose levels<br>(mg/dl) | 121.8±20.59 | 101.7±10.88 | 94.98±8.45 | <0.0001 | A-B<0.0001<br>A-C<0.0001<br>B-C=0.0623 |
| HbA1c (%) | 6.88±0.79 | - | - | - | - |
| Disease duration | 13.75±9.54 | - | - | - | - |
| Serum calcium (mg/dl) | 9.29± 0.43 | 9.20±0.44 | 9.16±0.34 | 0.659 | NS |
| eGFR(mL/min/1.73<br>m<sup>2</sup>) |  |  |  |  |  |
| Serum blood urea<br>nitrogen (mg/dl) | 45.40±14.23 | 45.07±13.36 | 38.52±9.10 | 0.076 | NS |
| Total cholesterol<br>(mg/dl) | 183.7±22.05 | 213.3±27.69 | 221.3±53.90 | 0.393 | NS |
| HDL cholesterol<br>(mg/dl) | 48.00±6.24 | 63.46±18.16 | 65.42±12.27 | 0.170 | NS |
| LDL cholesterol<br>(mg/dl) | 107.3±28.92 | 126.5±26.77 | 130.3±48.87 | 0.672 | NS |
| Triglycerides (mg/dl) | 165.0±35.16 | 123.0±48.45 | 111.3±50.32 | 0.208 | NS |

### Adipose tissue composition and bone mass

All the parameters related to adipose tissue composition and bone mass are reported in Table 2. Trunk total mass was significantly higher in subjects with obesity (p<0.0001) and T2D (p=0.001) compared to controls. Also, trunk fat mass was higher in OB (p=0.003) and T2D (p=0.016) compared to controls. Consistently, percent trunk fat was higher in subjects with obesity (p=0.001) and T2D (p=0.030) as compared to control subjects. Analysis of fat distribution highlighted that OB, but not T2D, had both a higher percentage of android fat (p<0.0001) and of gynoid fat (p=0.022) as compared to controls. Finally, levels of VAT mass and VAT volume were significantly greater in subjects with obesity (p<0.0001, respectively) and T2D (p=0.0002, respectively) compared to controls.

**Table 2.** Adipose tissue composition and bone mass by DXA Results were analyzed using unpaired Ordinary One-Way ANOVA and Tukey’s Multiple Comparisons test. Results are presented as mean ± standard deviation.

|  | **T2D (A)**<br>**n=19** | **OB (B)**<br>**n=17** | **CTRL (C)**<br>**N=27** | **P value** | **Multiple**<br>**Comparisons** |
| --- | --- | --- | --- | --- | --- |
| Trunk total mass (gr) | 39718±5094 | 42018±5861 | 31431±6753 | <0.0001 | A-C=0.001<br>B-C<0.0001 |
| Trunk fat mass (gr) | 19984±3637 | 20219±7444 | 14232±4870 | 0.0015 | A-C=0.016<br>B-C=0.003 |
| Trunk fat percentage (%) | 50.08±4.065 | 51.33±6.833 | 44.22±6.465 | 0.001 | A-C=0.030<br>B-C=0.001 |
| Android fat percentage (%) | 49.84±3.076 | 54.18±5.252 | 45.34±7.379 | 0.0001 | B-C<0.0001 |
| Gynoid fat percentage (%) | 48.52±5.356 | 50.62±4.704 | 46.17±5.349 | 0.028 | B-C=0.022 |
| VAT mass (gr) | 1170±284.8 | 1172±370.0 | 675.2±244.3 | <0.0001 | A-C=0.0002<br>B-C<0.0001 |
| VAT volume (cm<sup>3</sup>) | 1265±307.8 | 1267±400.1 | 729.8±264.3 | <0.0001 | A-C=0.0002<br>B-C<0.0001 |
| BMD total body (g/cm<sup>2</sup>) | 1.255±0.134 | 1.259±0.108 | 1.228±0.123 | 0.679 | NS |
| T score total body | 1.708±1.184 | 1.905±1.475 | 1.556±1.546 | 0.709 | NS |
| BMD Lumbar L1+L4 (g/cm<sup>2</sup>) | 0.9512±0.120 | 1.028±0.156 | 0.873±0.143 | 0.005 | B-C=0.004 |
| T score Lumbar | -0.866±1.094 | -0.166±1.422 | -1.581±1.305 | 0.005 | B-C=0.003 |
| BMD Femoral (g/cm<sup>2</sup>) | 0.856±0.125 | 0.873±0.119 | 0.727±0.070 | 0.0004 | A-C=0.016<br>B-C=0.0005 |
| T score Femoral | -0.685±1.042 | -0.550±0.976 | -1.854±0.543 | <0.0001 | A-C=0.006<br>B-C=0.0001 |

Total body bone mineral density (BMD) and T-score were not different between groups. Lumbar bone mineral density was higher in OB compared to controls (p=0.004) but not in T2D compared to controls. Lumbar T-score was also increased only in OB compared to controls (p=0.003). Femoral BMD was increased not only in OB (p=0.0005) but also in T2D (p= 0.016) compared to controls. In line, femoral T-score was higher in OB (p<0.0001) and T2D (p=0.006) compared to controls.

### Bone microarchitecture and strength

Microarchitecture parameters by μCT on trabecular core samples are reported in Table 3. T2D, OB and control subjects had similar trabecular indices, including bone volume density (BV/TV), connectivity density, trabecular thickness, trabecular number, trabecular separation, tissue mineral density and volumetric BMD. Biomechanic analysis (table 3) showed a reduced Young’s modulus in T2D compared to OB (p=0.0395) but not to control subjects, and a trend of reduction of Yield strength in T2D compared to OB (p=0.0552).

**Table 3.** Microarchitecture and mechanical parameters of trabecular bone of the study subjects, Results were analyzed using unpaired Ordinary One-Way ANOVA and Tukey’s Multiple Comparisons test. Results are presented as mean ± standard deviation.

|  | T2D (A)<br>n=19 | OB (B)<br>n=17 | CTRL (C)<br>N=18 | P value | Multiple<br>Comparisons |
| --- | --- | --- | --- | --- | --- |
| **Microcomputed tomography** |  |  |  |  |  |
| BV/TV | 0.240±0.135 | 0.236±0.096 | 0.291±0.092 | 0.396 | NS |
| Connectivity density(1/mm<sup>3</sup>) | 6.992±4.251 | 6.754±2.913 | 6.748±1.839 | 0.978 | NS |
| Trabecular number (1/mm) | 1.484±0.501 | 1.542±0.197 | 1.520±0.266 | 0.926 | NS |
| Trabecular thickness (mm) | 0.203±0.054 | 0.177±0.067 | 0.211±0.053 | 0.367 | NS |
| Trabecular separation (mm) | 0.709±0.197 | 0.585±0.114 | 0.582±0.090 | 0.074 | NS |
| Volumetric BMD (mgHA/cm<sup>3</sup>) | 220.4±100.2 | 224.1±78.07 | 263.5±68.33 | 0.381 | NS |
| Tissue mineral density (mgHA/cm<sup>3</sup>) | 689.4±32.80 | 680.7±29.59 | 690.7±29.80 | 0.681 | NS |
| **Biomechanical properties** |  |  |  |  |  |
| Young’s Modulus (MPa) | 23.68±10.82 | 69.28±59.23 | 50.75±46.06 | 0.050 | A-B=0.039 |
| Ultimate strength (MPa) | 3.632±2.178 | 5.607±3.344 | 5.199±3.021 | 0.256 | NS |
| Yield strength (MPa) | 2.410±1.160 | 4.849±2.777 | 4.591±2.442 | 0.049 | A-B=0.0559 |

### Circulating and bone inflammation

Circulating TNF-α levels were significantly higher in T2D (p=0.0004), but they were not higher in OB (p=0.1108) compared to control subjects (Fig. 1A). IL-6 levels were higher in both T2D (p=0.0451) and OB (p=0.0002) compared to controls (Fig. 1B). Adiponectin levels were lower only in T2D compared to OB (p=0.0402), but not in the other groups (Fig. 1C). Of note, circulating TNF-α and IL-6 were strongly positively correlated with VAT mass (r=0.3651, p=0.0310 and r=0.7139, p<0.0001, respectively), as reported in Figure 1-Supplementary figure 1.

**Figure 1.**
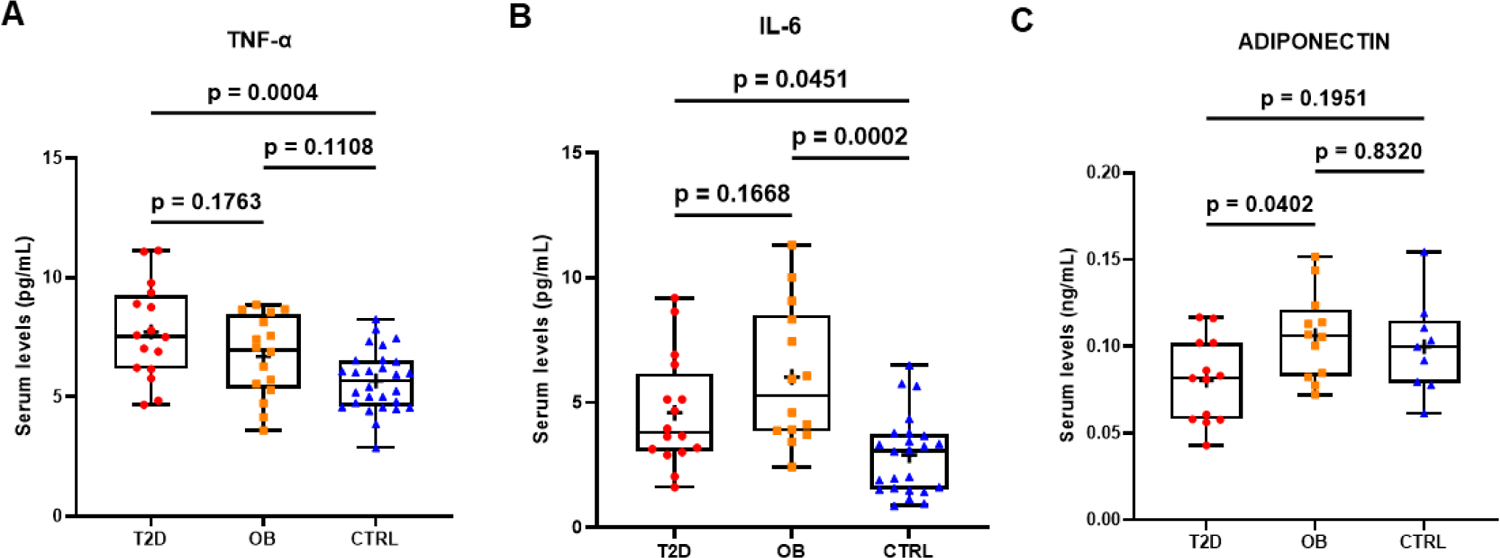
Circulating inflammatory cytokines of subjects with T2D, OB and controls. Serum (A) Tumor Necrosis Factor-alpha (TNF-α), (B) Interleukin-6 (IL-6), and (C) Adiponectin. The unit of TNF-α and IL-6 concentrations was pg/mL, while adiponectin concentration was ng/mL. The box plots show the medians (middle line) and first and third quartiles (boxes), and the whiskers show the minimum and maximum. Medians and interquartile ranges, differences between groups were analyzed using Ordinary One-Way ANOVA.

Analysis of gene expression in the femoral head after surgery showed increased TNF-α mRNA levels either in T2D compared to controls (p<0.0001) or in T2D compared to OB (p=0.0161) (Fig. 2A). However, gene expression levels of IL-6 were not different between groups (Fig. 2B), as for IL-8 mRNA levels (Fig. 2C) IL-10 gene expression levels were lower in both T2D (p=0.0285), and OB (p=0.0324) compared to controls (Fig. 2D). Adiponectin (ADIPOQ) mRNA levels were lower in subjects with T2D (p=0.0041) compared to controls (Fig. 2E). On the other hand, SFRP5 mRNA levels were only higher in T2D (p=0.0084) compared to control subjects (Fig. 2F).

**Figure 2.**
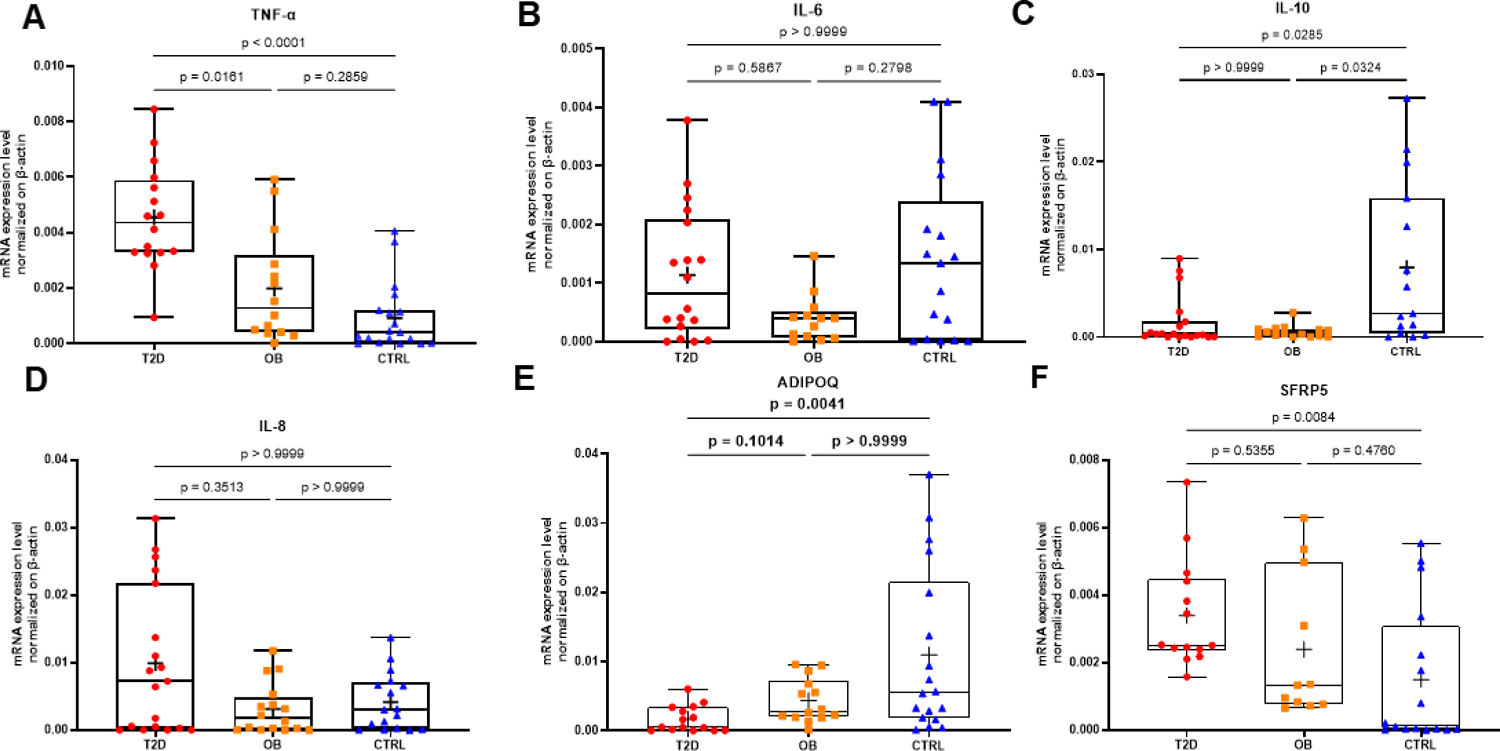
Gene expression analysis of inflammatory markers in trabecular bone samples of subjects with T2D, OB and controls. mRNA levels of (A) Tumor Necrosis Factor-alpha (TNF-α), (B) Interleukin-6 (IL-6), (C) Interleukin-10 (IL-10), (D) Interleukin-8 (IL-8), (E) Adiponectin (ADIPOQ), and (F) secreted frizzled-related protein 5 (SFRP5). Data are expressed as fold changes over beta-actin. Medians and interquartile ranges, differences between groups were analyzed using non-parametric One-Way ANOVA (Kruskal-Wallis test).

### Wnt signaling pathway in bone

The canonical Wnt inhibitor SOST mRNA levels were higher in T2D (p<0.0001) and in OB (p<0.0001) compared to controls (Fig.3A). Interestingly, SOST mRNA levels were positively correlated with fasting blood glucose levels in OB (Figure 3-Supplementary figure 1). In line, gene expression of the canonical ligand WNT10B was lower in T2D (p=0.0071) and in OB (p=0.0196) vs controls (Fig. 3B). Also, gene expression of LEF-1, the Wnt/β-catenin transcription factor, was lower in T2D (p<0.0001) and in OB (p=0.0479) compared to controls (Fig. 3C). Gene expression of non-canonical WNT5A was higher only in T2D (p=0.0025) but it was not different in OB compared to controls (Fig. 3D). Finally, Gene expression of GSK3β was higher either in T2D compared to OB (p=0.0069) or T2D vs controls (p=0.0003) (Fig. 3E).

**Figure 3.**
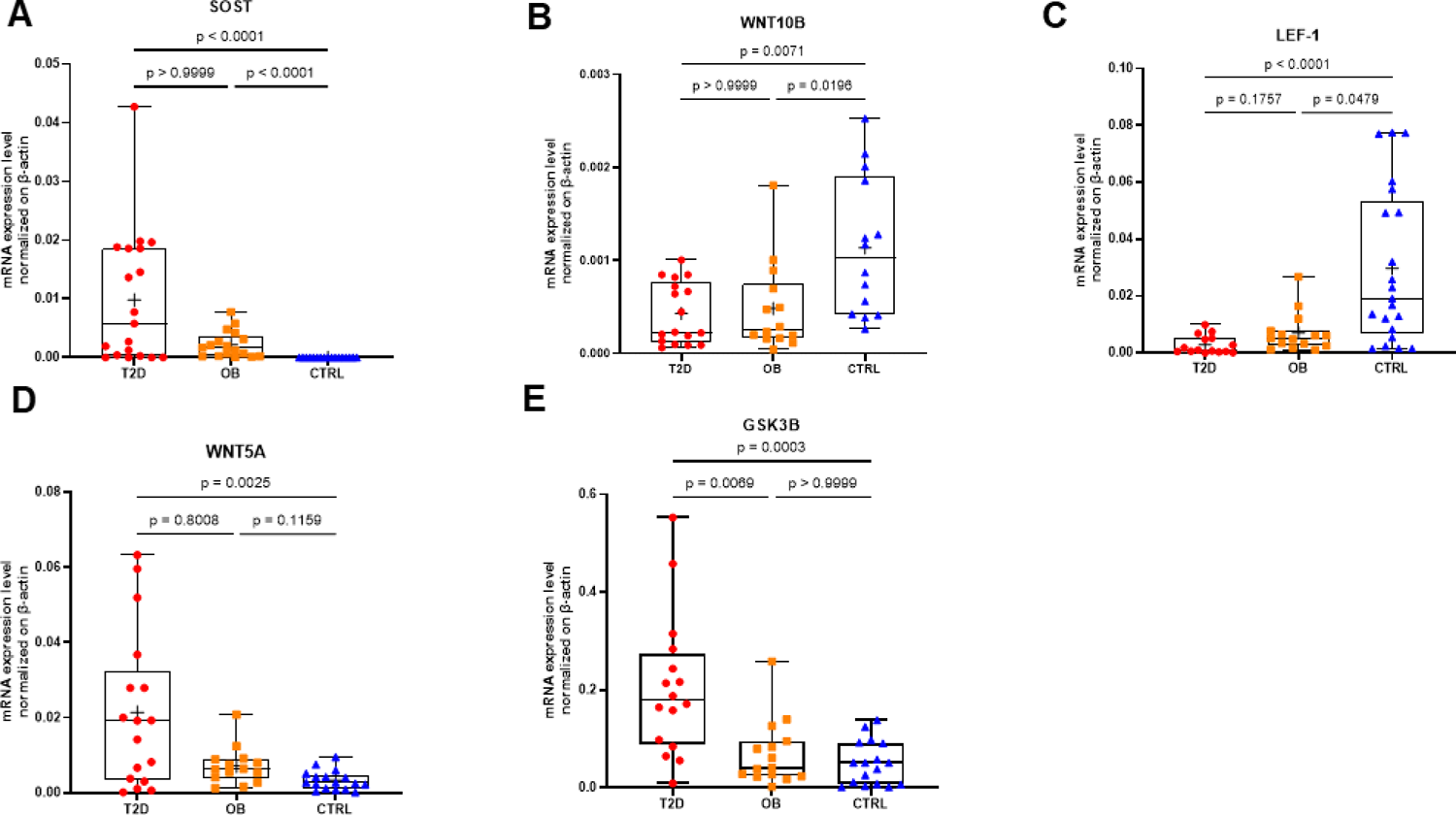
Gene expression analysis of Wnt signaling pathway in trabecular bone samples of subjects with T2D, OB and controls. mRNA levels of (A) Sclerostin (SOST) (B) Wnt family member 10B (WNT10B), (C) T-cell factor/ lymphoid enhancer factor 1 (LEF-1), (D) Wnt family member 5A (WNT5A), and (E) glycogen synthase kinase 3 beta (GSK3β). Data are expressed as fold changes over beta-actin. Medians and interquartile ranges, differences between groups were analyzed using non-parametric One-Way ANOVA (Kruskal-Wallis test).

### Correlation analysis of inflammatory and Wnt target genes

Correlation analysis of inflammatory and Wnt target genes are reported in figure 4 (A-I). TNF-α mRNA levels were positively correlated with SOST (r=0.5121, p=0.0002), WNT5A (r=0.3227, p=0.0396) and GSK3β (r=0.3789, p=0.0146) mRNA levels (Fig. 4A-C). Conversely, TNF-α was negatively correlated with WNT10B (r=-0.3844, p=0.0188) and LEF-1 (r=-0.3310, p=0.0322) mRNA levels (Fig. 4D,E). IL-10 was only negatively correlated with SOST (r=-0.3100, p=0.0457) (Fig. 4F). ADIPOQ mRNA levels were negatively correlated with SOST (r=-0.3864, p=0.0105) and with WNT5A mRNA levels (r=-0.3025, p=0.0515) (Fig. 4G,H). Moreover, SFRP5 mRNA levels negatively correlated with LEF-1 (r=-0.3991, p=0.0131) mRNA levels (Fig. 4I).

**Figure 4.**
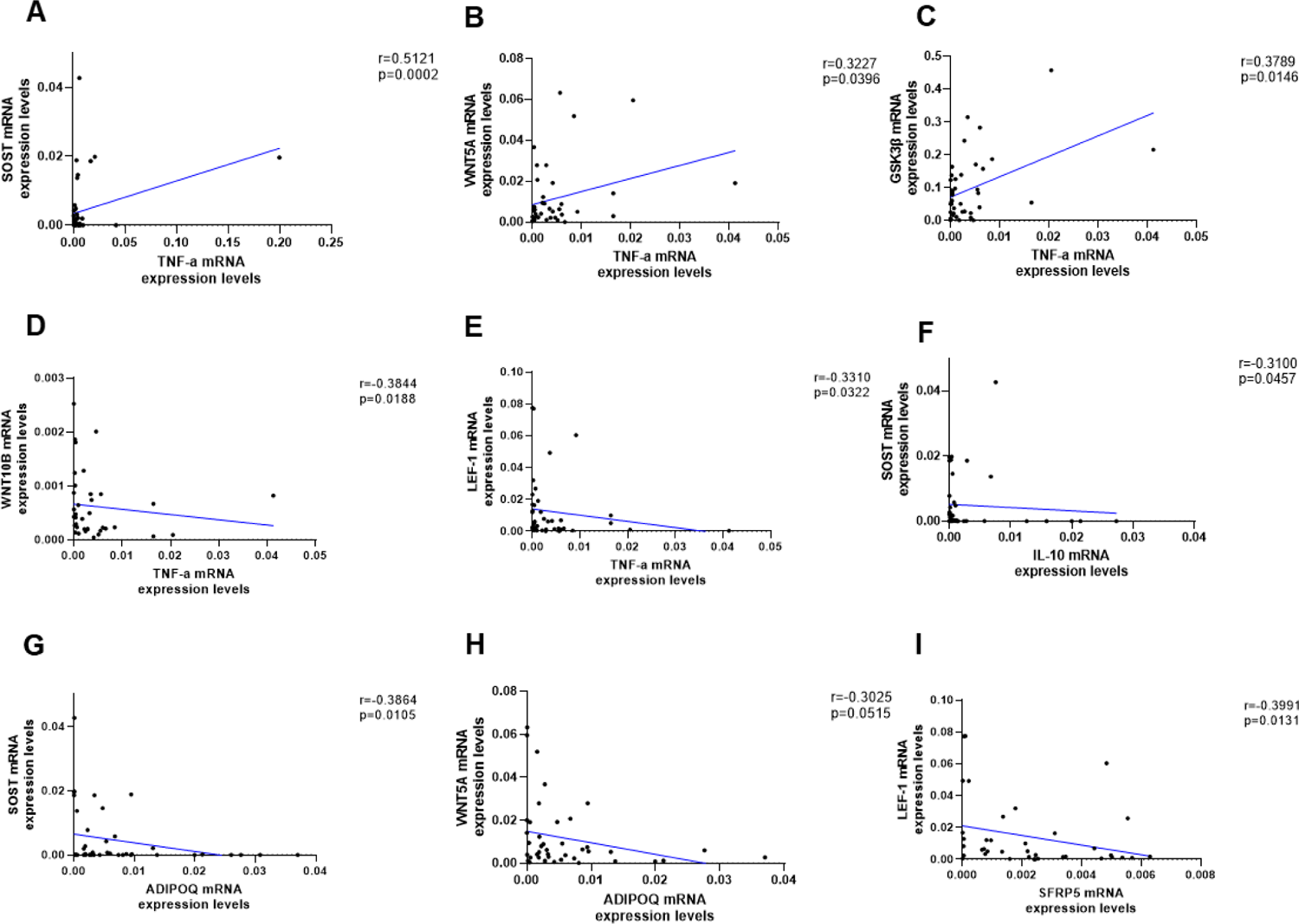
Correlation analysis of inflammatory and Wnt target genes in all study subjects. Correlations between Tumor Necrosis Factor-alpha (TNF-α) and (A) sclerostin (SOST), (B) Wnt family member 5A (WNT5A), (C) glycogen synthase kinase 3 beta (GSK3β), (D) Wnt family member 10B (WNT10B), and (E) T-cell factor/ lymphoid enhancer factor 1 (LEF-1). (F) Correlation between Interleukin-10 (IL-10) and sclerostin (SOST). Correlation between Adiponectin (ADIPOQ) and (G) sclerostin (SOST), (H) Wnt family member 5A (WNT5A). (I) Correlation between secreted frizzled-related protein 5 (SFRP5) and T-cell factor/ lymphoid enhancer factor 1 (LEF-1). Data were analyzed using nonparametric Spearman correlation analysis and r represents the correlation coefficient.

### Correlation analysis of inflammatory cytokines and bone strength

Serum levels of TNF-α (r=-0.3473, p=0.0352) and IL-6 (r=-0.3777, p=0.0302) negatively correlated with Young’s Modulus (Fig. 5A-B). There was no difference between circulating inflammatory markers and the other biomechanical parameters.

**Figure 5.**
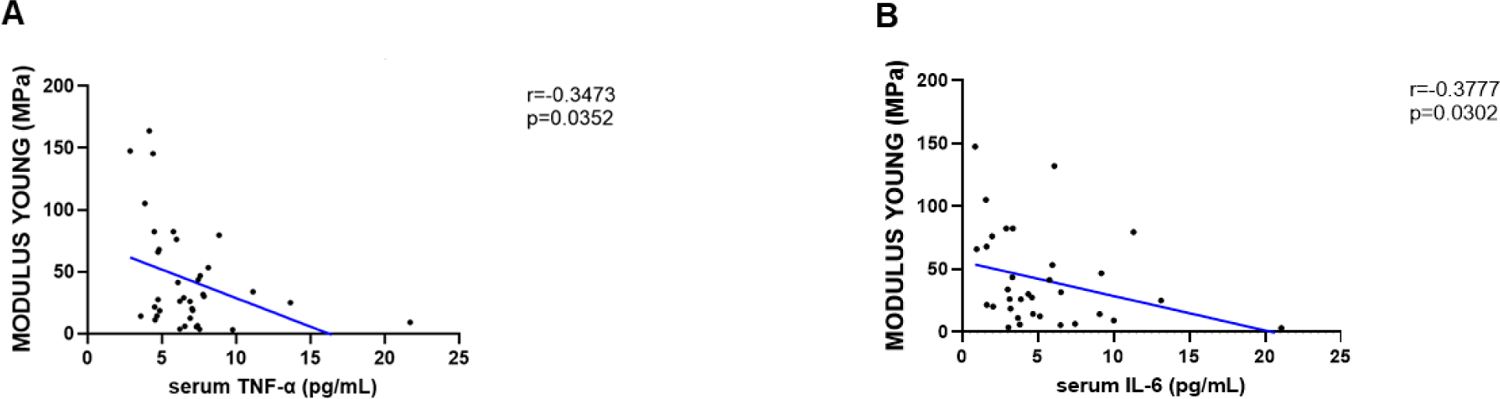
Correlation analysis of circulating inflammatory cytokines and Young’s Modulus in all study subjects. Correlations between (A) Tumor Necrosis Factor-alpha (TNF-α) and Young’s Modulus, and (B) Interleukin-6 (IL-6) and Young’s Modulus. Data were analyzed using nonparametric Spearman correlation analysis and r represents the correlation coefficient.

## Discussion

In this study we showed increased circulating and bone inflammation in postmenopausal women with T2D and obesity. Specifically, we found increased TNF-α mRNA levels in bone of subjects with T2D while the anti-inflammatory cytokine IL-10 was decreased in both subjects with T2D and obesity. Importantly, increased mRNA levels of inflammatory cytokines were associated with downregulation of Wnt pathway related genes. Moreover, serum TNF-α was increased in T2D compared to the other groups and it was negatively associated with bone strength.

The increased TNF-α serum levels in subjects with T2D observed in our study are consistent with the study of Sum et al. [3] that reported increased TNF-α and other chemokines in T2D patients with bone fracture. There are evidence showing that inflammation may affect bone health. *In vitro* studies showed that TNF-α is involved in the regulation of bone resorption in inflammatory conditions either directly or by increasing RANKL and M-CSF expressions by osteoblasts, stromal cells and even osteocytes [15]. Moreover, blocking TNF-α in bone-lining forming cells increased cell number and enhanced bone formation in a model of T2D [10]. In line, we found that both serum and bone TNF-α were increased in T2D, but not in subjects with obesity. Interestingly, serum TNF-α was negatively correlated with Young’s Modulus, suggesting that increased TNF-α reduces bone strength in subjects with T2D.

Although we confirmed that circulating IL-6 was higher in subject with obesity and it was positively correlated with VAT, we found no difference of IL-6 gene expression levels in bone tissue of either T2D or subjects with obesity. There is only evidence that IL-6 contributed to bone loss *in vitro*, affecting BMCs differentiation and cellular senescence in a model of high fat diet-induced obesity [9]. However, there is no data supporting the effect of inflammation in bone of subjects with obesity. In line with IL-6, gene expression of pro-inflammatory IL-8 was not altered either in bone of subjects with T2D or obesity. There is no evidence about IL-8 role in bone of either T2D or obesity [8].

On the other hand, we found that the anti-inflammatory IL-10 mRNA levels were decreased in subjects with T2D and obesity. Previous data showed that serum levels of IL-10 were substantially lower in osteoporosis patients [16] and in T2D [17] than in healthy individuals, although to our knowledge, the role of IL-10 on bone of T2D or obesity has not been studied. As expected, both serum and mRNA bone adiponectin levels were lower in T2D [18,19]. *In vitro* studies indicated that adiponectin promoted osteoblastogenesis, whilst simultaneously inhibiting osteoclastogenesis [20–22] and adiponectin knockout mice have increased bone mass and osteoblast numbers [23].

Moreover, serum adiponectin in postmenopausal women was found inversely associated with bone mass [24]. However, there are no data about adiponectin in bone of subjects with T2D or obesity. We also showed that subjects with T2D had increased SFRP5 bone mRNA levels compared to controls. Data reported that SFRP5, a negative regulator of Wnt canonical signaling pathway, was overexpressed in osteoporosis and was also associated to downregulated bone formation markers [25]. Our data are in line with these findings, supporting that T2D is characterized by low bone formation and downregulated Wnt canonical signaling.

Previous studies showed that inflammation may downregulate Wnt signaling pathway [26], the main regulator of bone metabolism. Consistent with our previous findings [14], we observed that the expression of the Wnt inhibitor SOST was not only higher in bone of T2D but also in subjects with obesity compared to controls. Even though sclerostin is increased during weight loss [27], there are other data showing that serum sclerostin was negatively associated to insulin sensitivity in women with obesity [28]. Conversely, our data showed that SOST mRNA levels were positively correlated with fasting blood glucose in subjects with obesity suggesting that glucose impairment may be a determinant for higher sclerostin levels. Consistent with increased SOST mRNA levels and in line with our previous findings on Wnt signaling regulation in bone of T2D [14], the expression of the Wnt ligand WNT10B and the transcription factor LEF-1 were downregulated in both subjects with T2D and obesity. Also, *in vitro* studies showed that TNF-α induced the phosphorylation of GSK3β, the deactivated form of GSK3β, which increased nuclear β-catenin and Lef-1 resulting in downregulated Wnt signaling and reduced bone formation [26].Our data confirmed that GSK3βB is increased in subjects with T2D. This is in line with our previous findings in which GSK3β was associated with reduced bone formation and strength in T2D [14].

Furthermore, it is shown that Wnt5a promotes inflammation and insulin resistance in adipose tissue of human and animal models [29,30] and high circulating levels of Wnt5a are detected in subjects with T2D [31] and obesity [32]. Other studies showed that Wnt5a inhibits Wnt3a protein by downregulating beta/catenin-induced reporter gene expression [33]. In line, we observed that gene expression of WNT5A was increased in bone of T2D. This study has some limitations. It is a cross-sectional design with a relatively small number of enrolled subjects.

The main strength of the study is that we investigated for the first time the effect of inflammation on human bone samples measuring several pro and anti-inflammatory genes. Moreover, we found the association between inflammation and bone strength by the regulation of Wnt signaling pathway in subjects with T2D and obesity compared to a control population.

In conclusion, our study showed that there is increased inflammation in bone of subjects with T2D and obesity and it is associated with the downregulation of Wnt signaling and reduced bone strength. These results shed light on the pathophysiology of bone impairment in T2D and obesity and may help understanding the mechanisms of bone fragility in metabolic diseases.

## Material and methods

### Study subjects

We enrolled for this study 63 postmenopausal women (19 with T2D, 17 with obesity and 27 non-diabetic normal weight controls) undergoing hip replacement surgery for osteoarthritis, consecutively screened to participate in this study between 2018 and 2022. Diabetes status was confirmed by the diabetologist. Participants were diagnosed with type 2 diabetes when they had fasting plasma glucose (FPG) ≥126 mg/dl or 2-h plasma glucose (2-h PG) ≥200 mg/dl during a 75-g oral glucose tolerance test (OGTT); or hemoglobin A1c (HbA1c) ≥6.5% in accordance with the American Diabetes Association diagnostic criteria [34]. Obesity was confirmed based on the BMI ≥30 kg/m^2^, in accordance with the World Health Organization diagnostic criteria [35]. Eligible participants were ≥60 years of age. Exclusion criteria were any diseases affecting bone, any concomitant inflammatory disease or malignancy. Additionally, individuals treated with medications affecting bone metabolism such as estrogen, raloxifene, tamoxifen, bisphosphonates, teriparatide, denosumab, thiazolidinediones, glucocorticoids, anabolic steroids, and phenytoin, and those with hypercalcemia or hypocalcemia, hepatic or renal disorder, hypercortisolism, current alcohol or tobacco use were excluded. The study was approved by the Ethics Committee of Campus Bio-Medico University of Rome (Prot..42/14 PT_ComEt CBM) and all participants provided written informed consent. All procedures were conducted in accordance with the Declaration of Helsinki.

### Body Composition/Anthropometric measurements

Body scans were conducted with Lunar ProdigyTM (GE Healthcare©, USA) DXA scanner. Body composition parameters were analyzed using en-CORETM software (version 17, 2016), and VAT was measured with the CoreScanTM (GE Healthcare©, USA). Quality control and calibration of the model were performed each day, following the instruction protocol provided by the manufacturer. For this analysis, the variables of interest derived from the DXA dataset are BMD total body (g/cm^2^), T score total body, BMD Lumbar L1+L4 (g/cm^2^), T score femoral, trunk fat mass (gr), trunk total mass (gr), and defined anatomical regions (android, and gynoid % fat) and visceral adipose tissue (VAT mass, g and VAT volume, cm^3^). The body mass index (BMI) was calculated as weight/(height)^2^ (kg/m^2^).

### Processing of blood Samples

Venous whole blood was collected the day before the surgery in all participants after completing a fasting period of at least 8 hours, using tubes containing clot activator (BD vacutainer®, NJ, USA). Samples were kept at room temperature to allow the blood to clot, and they were centrifuged at 1500 g for 10 minutes for serum isolation. Serum was then transferred to a clean tube and stored at −80°C for further analysis.

### Processing of bone samples

Femoral head specimens were obtained during hip replacement surgery. Briefly, trabecular bone specimens were collected fresh and washed multiple times in sterile PBS until the supernatant was clear of blood, as previously described [13]. Trabecular samples were then aliquoted and stored at −80°C for further analysis.

### ELISA and ELLA Microfluidic Immunoassay

Adiponectin was quantified in serum using ELISA immunoassay (R&D system, MN, USA) according to the manufacturer instructions. TNF-α and IL-6 were measured in serum by using Simple Plex assays run on the ELLA microfluidic immunoassay system (ProteinSimple, San Jose, CA). 25 μl of serum was diluted at a 1:1 ratio with sample diluent, and 50 μl the solution was added to each sample inlet on the ELLA cartridge, according to manufacturer’s instruction. Sample results were reported using Simple Plex Runner v.3.7.2.0 (ProteinSimple).

### RNA extraction and gene expression by RT-PCR

Total RNA from trabecular bone samples was extracted using TRIzol (Invitrogen) following the manufacturer’s instructions. The concentration and purity of the extracted RNA were assessed spectrophotometrically (TECAN, InfiniteM200PRO), and only samples with 260/280 absorbance ratio between 1.8 and 2 were used for reverse transcription using High-Capacity cDNA Reverse Transcription Kit (Applied Biosystems, Carlsbad, CA) according to the manufacturer’s recommendations. Transcription products were amplified using TaqMan real-time PCR (Applied Biosystems, Carlsbad, CA) and a standard protocol (95°C for 10 minutes; 40 cycles of 95°C for 15 seconds and 60°C for 1 minute; followed by 95°C for 15 seconds, 60°C for 15 seconds, and 95°C for 15 seconds). β-Actin expression was used as an internal control (housekeeping gene). Relative expression levels of Tumor necrosis factor alpha (TNF-α), Interleukin 6 (IL-6), Adiponectin (ADIPOQ), Interleukin 8 (IL-8), Interleukin 10 (IL-10), secreted frizzled-related protein 5 (SFRP5), Sclerostin (SOST), Wnt ligands (WNT5A and WNT10B), T-cell factor/ lymphoid enhancer factor 1 (LEF-1), and glycogen synthase kinase 3 beta (GSK3B), were calculated using the 2-ΔCt method.

### Micro-computed tomography (μCT) and bone compression tests

We used cylindrical bone specimens of trabecular core (with a diameter of 10 mm and a length of 20 mm) from 19 T2D with obesity, 17 OB non-diabetic and 18 control subjects to measure bone micro-architecture and bone mechanical parameters (Young’s modulus, ultimate strength and yield strength), as previously described [13].

### Statistical analysis

Data were analyzed using GraphPad Prism 9.0 (GraphPad Software, San Diego, CA). Patients’ characteristics were described using means and standard deviations (SD) or medians and 25th-75th percentiles, as appropriate, and percentages. Group data are presented in boxplots with median and interquartile range (IQR); whiskers represent maximum and minimum values. We assessed data for normality and Mann-Whitney test was used to compare variables between groups. Data were analyzed using nonparametric Spearman correlation analysis and the correlation coefficients (r) were used to assess the relationship between variables. We used Grubbs’ Test to assess and exclude outliers.

## Acknowledgements

This work was supported by an internal Grant of Campus Bio-Medico University of Rome. We wish to thank the structure & strength core at Washington University Musculoskeletal Research Center.

## Supplementary figure legends

**Figure 1-Supplementary figure 1.**
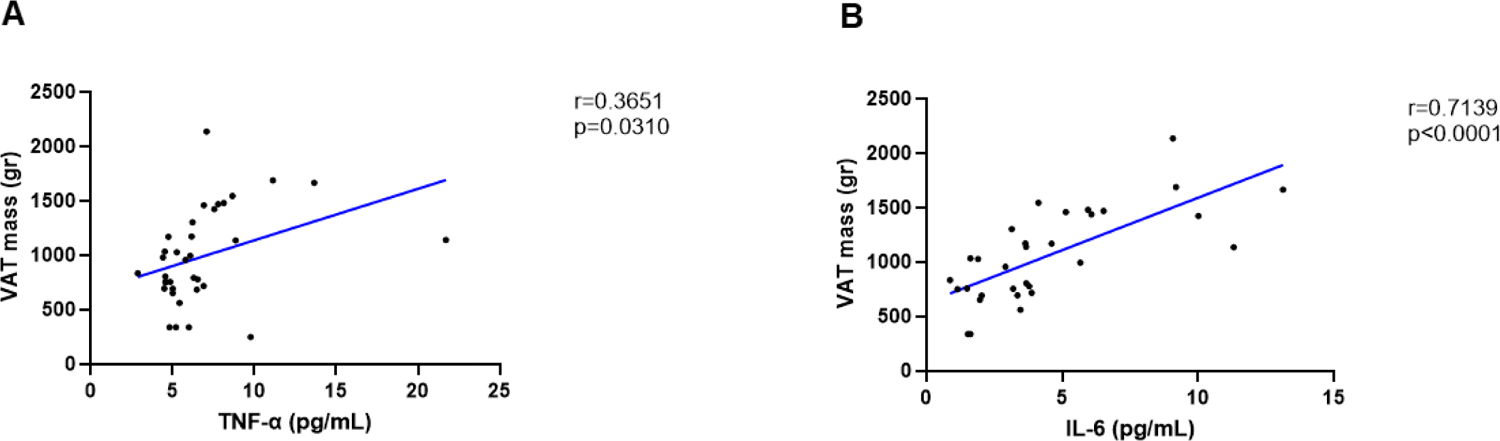
Correlation analysis of circulating inflammatory cytokines and Visceral adipose tissue (VAT) mass in all study subjects. Correlations between (A) Tumor Necrosis Factor-alpha (TNF-α) and VAT mass, and (B) Interleukin-6 (IL-6) and VAT mass. Data were analyzed using Pearson correlation analysis and r represents the correlation coefficient.

**Figure 3-Supplementary figure 1.**
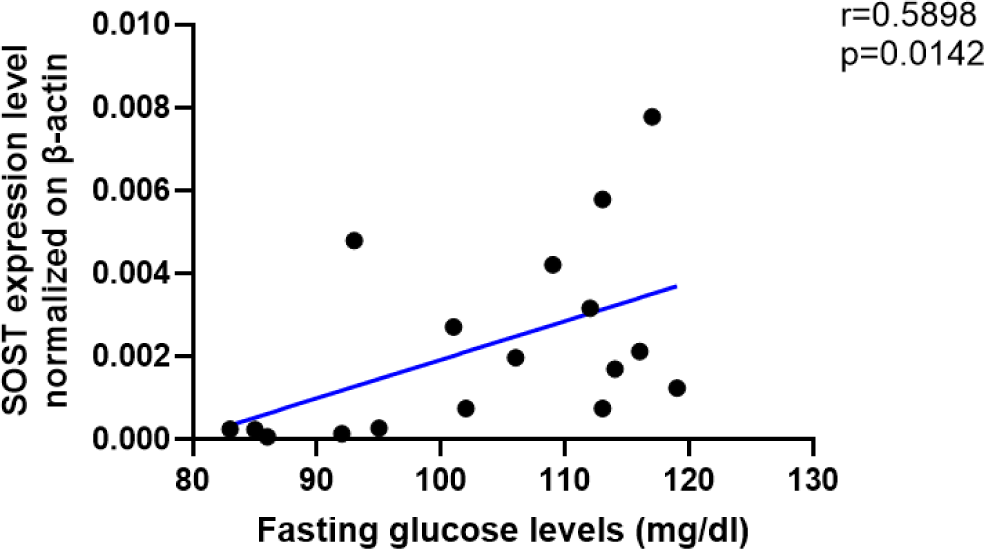
Correlation analysis of SOST mRNA levels in bone and fasting blood glucose in subjects with obesity (OB). Data were analyzed using nonparametric Spearman correlation analysis and r represents the correlation coefficient.

## References

[1] Rohm T V, Meier DT, Olefsky JM, Donath MY. Inflammation in obesity, diabetes, and related disorders. Immunity 2022;55:31–55. 10.1016/j.immuni.2021.12.013.

[2] Park HS, Park JY, Yu R. Relationship of obesity and visceral adiposity with serum concentrations of CRP, TNF-alpha and IL-6. Diabetes Res Clin Pract 2005;69:29–35. 10.1016/j.diabres.2004.11.007.

[3] Sun M, Yang J, Wang J, Hao T, Jiang D, Bao G, et al. TNF-α is upregulated in T2DM patients with fracture and promotes the apoptosis of osteoblast cells in vitro in the presence of high glucose. Cytokine 2016;80:35–42. 10.1016/j.cyto.2016.01.011.

[4] Bathina S, Armamento-Villareal R. The complex pathophysiology of bone fragility in obesity and type 2 diabetes mellitus: therapeutic targets to promote osteogenesis. Front Endocrinol (Lausanne) 2023;14:1168687. 10.3389/fendo.2023.1168687.

[5] Cauley JA, Barbour KE, Harrison SL, Cloonan YK, Danielson ME, Ensrud KE, et al. Inflammatory Markers and the Risk of Hip and Vertebral Fractures in Men: the Osteoporotic Fractures in Men (MrOS). J Bone Miner Res 2016;31:2129–38. 10.1002/jbmr.2905.

[6] Barbour KE, Boudreau R, Danielson ME, Youk AO, Wactawski-Wende J, Greep NC, et al. Inflammatory markers and the risk of hip fracture: the Women’s Health Initiative. J Bone Miner Res 2012;27:1167–76. 10.1002/jbmr.1559.

[7] Barbour KE, Lui L-Y, Ensrud KE, Hillier TA, LeBlanc ES, Ing SW, et al. Inflammatory markers and risk of hip fracture in older white women: the study of osteoporotic fractures. J Bone Miner Res 2014;29:2057–64. 10.1002/jbmr.2245.

[8] Xu J, Yu L, Liu F, Wan L, Deng Z. The effect of cytokines on osteoblasts and osteoclasts in bone remodeling in osteoporosis: a review. Front Immunol 2023;14:1222129. 10.3389/fimmu.2023.1222129.

[9] Li Y, Lu L, Xie Y, Chen X, Tian L, Liang Y, et al. Interleukin-6 Knockout Inhibits Senescence of Bone Mesenchymal Stem Cells in High-Fat Diet-Induced Bone Loss. Front Endocrinol (Lausanne) 2020;11:622950. 10.3389/fendo.2020.622950.

[10] Liu R, Bal HS, Desta T, Behl Y, Graves DT. Tumor necrosis factor-alpha mediates diabetes-enhanced apoptosis of matrix-producing cells and impairs diabetic healing. Am J Pathol 2006;168:757–64. 10.2353/ajpath.2006.050907.

[11] Zhao B. TNF and Bone Remodeling. Curr Osteoporos Rep 2017;15:126–34. 10.1007/s11914-017-0358-z.

[12] Diarra D, Stolina M, Polzer K, Zwerina J, Ominsky MS, Dwyer D, et al. Dickkopf-1 is a master regulator of joint remodeling. Nat Med 2007;13:156–63. 10.1038/nm1538.

[13] Piccoli A, Cannata F, Strollo R, Pedone C, Leanza G, Russo F, et al. Sclerostin Regulation, Microarchitecture, and Advanced Glycation End-Products in the Bone of Elderly Women With Type 2 Diabetes. J Bone Miner Res 2020;35:2415–22. 10.1002/jbmr.4153.

[14] Leanza G, Cannata F, Faraj M, Pedone C, Viola V, Tramontana F, et al. Bone canonical Wnt signaling is downregulated in type 2 diabetes and associates with higher advanced glycation end-products (AGEs) content and reduced bone strength. Elife 2024;12. 10.7554/eLife.90437.

[15] Marahleh A, Kitaura H, Ohori F, Kishikawa A, Ogawa S, Shen W-R, et al. TNF-α Directly Enhances Osteocyte RANKL Expression and Promotes Osteoclast Formation. Front Immunol 2019;10:2925. 10.3389/fimmu.2019.02925.

[16] Ma X, Zhu X, He X, Yi X, Jin A. The Wnt pathway regulator expression levels and their relationship to bone metabolism in thoracolumbar osteoporotic vertebral compression fracture patients. Am J Transl Res 2021;13:4812–8.

[17] van Exel E, Gussekloo J, de Craen AJM, Frölich M, Bootsma-Van Der Wiel A, Westendorp RGJ, et al. Low production capacity of interleukin-10 associates with the metabolic syndrome and type 2 diabetes: the Leiden 85-Plus Study. Diabetes 2002;51:1088–92. 10.2337/diabetes.51.4.1088.

[18] Hotta K, Funahashi T, Arita Y, Takahashi M, Matsuda M, Okamoto Y, et al. Plasma concentrations of a novel, adipose-specific protein, adiponectin, in type 2 diabetic patients. Arterioscler Thromb Vasc Biol 2000;20:1595–9. 10.1161/01.atv.20.6.1595.

[19] Jalovaara K, Santaniemi M, Timonen M, Jokelainen J, Kesäniemi YA, Ukkola O, et al. Low serum adiponectin level as a predictor of impaired glucose regulation and type 2 diabetes mellitus in a middle-aged Finnish population. Metabolism 2008;57:1130–4. 10.1016/j.metabol.2008.03.019.

[20] Lee HW, Kim SY, Kim AY, Lee EJ, Choi J-Y, Kim JB. Adiponectin stimulates osteoblast differentiation through induction of COX2 in mesenchymal progenitor cells. Stem Cells 2009;27:2254–62. 10.1002/stem.144.

[21] Lin YY, Chen CY, Chuang TY, Lin Y, Liu HY, Mersmann HJ, et al. Adiponectin receptor 1 regulates bone formation and osteoblast differentiation by GSK-3β/β-catenin signaling in mice. Bone 2014;64:147–54. 10.1016/j.bone.2014.03.051.

[22] Wu X, Huang L, Liu J. Effects of adiponectin on osteoclastogenesis from mouse bone marrow-derived monocytes. Exp Ther Med 2019;17:1228–33. 10.3892/etm.2018.7069.

[23] Kajimura D, Lee HW, Riley KJ, Arteaga-Solis E, Ferron M, Zhou B, et al. Adiponectin regulates bone mass via opposite central and peripheral mechanisms through FoxO1. Cell Metab 2013;17:901–15. 10.1016/j.cmet.2013.04.009.

[24] Napoli N, Pedone C, Pozzilli P, Lauretani F, Ferrucci L, Incalzi RA. Adiponectin and bone mass density: The InCHIANTI study. Bone 2010;47:1001–5. 10.1016/j.bone.2010.08.010.

[25] Chen H, He Y, Wu D, Dai G, Zhao C, Huang W, et al. Bone marrow sFRP5 level is negatively associated with bone formation markers. Osteoporos Int 2017;28:1305–11. 10.1007/s00198-016-3873-3.

[26] Kong X, Liu Y, Ye R, Zhu B, Zhu Y, Liu X, et al. GSK3β is a checkpoint for TNF-α-mediated impaired osteogenic differentiation of mesenchymal stem cells in inflammatory microenvironments. Biochim Biophys Acta 2013;1830:5119–29. 10.1016/j.bbagen.2013.07.027.

[27] Armamento-Villareal R, Sadler C, Napoli N, Shah K, Chode S, Sinacore DR, et al. Weight loss in obese older adults increases serum sclerostin and impairs hip geometry but both are prevented by exercise training. J Bone Miner Res 2012;27:1215–21. 10.1002/jbmr.1560.

[28] Aznou A, Meijer R, van Raalte D, den Heijer M, Heijboer A, de Jongh R. Serum sclerostin is negatively associated with insulin sensitivity in obese but not lean women. Endocr Connect 2021;10:131–8. 10.1530/EC-20-0535.

[29] Fuster JJ, Zuriaga MA, Ngo DT-M, Farb MG, Aprahamian T, Yamaguchi TP, et al. Noncanonical Wnt signaling promotes obesity-induced adipose tissue inflammation and metabolic dysfunction independent of adipose tissue expansion. Diabetes 2015;64:1235–48. 10.2337/db14-1164.

[30] Zuriaga MA, Fuster JJ, Farb MG, MacLauchlan S, Bretón-Romero R, Karki S, et al. Activation of non-canonical WNT signaling in human visceral adipose tissue contributes to local and systemic inflammation. Sci Rep 2017;7:17326. 10.1038/s41598-017-17509-5.

[31] Lu Y-C, Wang C-P, Hsu C-C, Chiu C-A, Yu T-H, Hung W-C, et al. Circulating secreted frizzled-related protein 5 (Sfrp5) and wingless-type MMTV integration site family member 5a (Wnt5a) levels in patients with type 2 diabetes mellitus. Diabetes Metab Res Rev 2013;29:551–6. 10.1002/dmrr.2426.

[32] Schulte DM, Müller N, Neumann K, Oberhäuser F, Faust M, Güdelhöfer H, et al. Pro-inflammatory wnt5a and anti-inflammatory sFRP5 are differentially regulated by nutritional factors in obese human subjects. PLoS One 2012;7:e32437. 10.1371/journal.pone.0032437.

[33] Mikels AJ, Nusse R. Purified Wnt5a protein activates or inhibits beta-catenin-TCF signaling depending on receptor context. PLoS Biol 2006;4:e115. 10.1371/journal.pbio.0040115.

[34] American Diabetes Association. (2) Classification and diagnosis of diabetes. Diabetes Care 2015;38 Suppl:S8–16. 10.2337/dc15-S005.

[35] who BMI n.d. https://www.who.int/health-topics/obesity#tab=tab_1.

